# Global increase in the endemism of birds from north to south

**DOI:** 10.1101/2024.05.30.596746

**Authors:** D. Matthias Dehling, Steven L. Chown

## Abstract

Endemism is a highly valuable metric for conservation because it identifies areas with irreplaceable species, ecological functions, or evolutionary lineages^1–6^. Global analyses of endemism currently fail to identify the most irreplaceable areas because the commonly used endemism metrics are correlated with richness, and entire regions, especially in the southern hemisphere, are regularly excluded^7–11^. Global patterns of endemism are therefore still insufficiently known. Here, using metrics representing irreplaceability, we unveil global patterns of avian taxonomic, functional, and phylogenetic endemism that show striking differences between hemispheres. Across all facets of diversity, endemism decreases poleward in the northern, but increases poleward in the southern, hemisphere, resulting in a global north-south increase in endemism. The pattern is driven by increasingly smaller and discontinuous landmasses towards the south leading to increasingly smaller ranges and reduced overlap in community composition, and, unexpectedly, to peaks of diversity relative to available area in the southern hemisphere. The current misapprehension of global endemism potentially compromises urgent conservation actions, drawing attention away from key areas of irreplaceability. Highly endemic southern-hemisphere communities might be especially vulnerable to the climate crisis because discontinuous landmasses impede range shifts.

---

Endemism serves as a measure for the irreplaceability of species because it describes species that occur only in a predefined area (e.g., habitat or country^1–3^) or, more generally, species with restricted ranges^4–6^. Analyses of endemism are a crucial component of global-scale conservation assessments because they identify areas of globally unique biodiversity that are complementary to areas rich in species^2,12–15^. Together they form the foundation of understanding global variation in diversity and the efforts required to conserve it^12,16^.

Current assessments of global endemism are, however, severely biased. Global diversity studies routinely exclude species-poor areas, especially in the southern hemisphere, and most notably the Antarctic region^8–11^. While the omission of sites does not affect estimates of alpha diversity (local species richness) in the remaining sites, it invariably alters estimates of endemism because endemism is assessed by comparing the composition of an assemblage with the composition of all other assemblages. In addition, the most commonly used measure for endemism (weighted endemism) is strongly influenced by local species richness^4,7^. This measure is therefore limited in its suitability for identifying sites with the highest concentration of range-restricted species. That is, the sites whose composition differs most from those of all other sites, and which are therefore most valuable for conservation because they harbour the highest concentration of unique species, functional trait combinations or phylogenetic lineages. Current assessments of global endemism potentially misrepresent such areas of high endemism, but the extent of this bias is unknown.

Moreover, if endemism is calculated based on biased evidence and then used to assess the biodiversity conservation value of sites and guide implementation decisions—as is routinely done at continental and global scales^7,15,17–19^—, implementation may fail to deliver the intended conservation outcomes. Doing so may instead inadvertently draw attention away from those sites whose communities are most dependent on range-restricted species and which are therefore most vulnerable to conservation threats^20,21^. These threats include those such as species invasions^22^, and/or climate change potentially shifting species ranges outside conservation areas^23–24^ or making current ranges unsuitable^25–27^. Despite the urgency of implementing conservation measures, given rapidly developing threats and declines in biodiversity^21,28^, the global distribution of irreplaceable diversity is still insufficiently known^29^.

To address these substantial deficiencies, we analysed global patterns of endemism and diversity of birds for the three key facets of diversity^30^: taxonomic diversity (i.e. richness), functional diversity, and phylogenetic diversity, across all global areas using complementarity (also known as *corrected weighted endemism*^4,31^ or *proportional range rarity*^32^) as a metric^33–36^. Complementarity measures how much of a site’s diversity (species, functional trait combinations, or phylogenetic lineages) is shared with other sites; it is independent of richness, and it peaks at sites whose composition differs most from all other sites, thereby providing an accurate picture of endemism as irreplaceability. Complementarity has rarely been analysed on global scales^29,32^, and less so for all facets of diversity combined.

Specifically (see details in Methods section), we compiled the most recent global datasets on avian breeding ranges^37^, functional traits^38^, and phylogeny^39^, and projected range maps onto a grid of 14,640 hexagonal grid cells (size: 11,610 km²) in an equal-area projection (EPSG:6933). For each grid cell, we determined the presence/absence of each bird species, functional trait combination, and phylogenetic lineage and then calculated alpha diversity (local richness), contribution to gamma diversity (i.e. “weighted endemism”), and complementarity. Since contribution to gamma and complementarity are both influenced by the overlap in assemblage composition and, hence, species’ range size, we also determined absolute range size, latitudinal range, longitudinal range, and latitudinal range midpoint for each bird species, functional trait combination, and branch in the phylogeny. For each measure, we calculated median values across latitudinal bands of 5° latitude and then analysed the relationship with latitude using linear models.

Alpha diversity peaks in the tropics and generally declines towards higher latitudes in both hemispheres (Fig. 1a,b). It thus follows the classic pattern of a latitudinal gradient in diversity^40^, albeit the relationships are slightly bi-modal as alluded to previously^41–42^, with a local minimum at about 22.5 °—more prominently shown in the northern hemisphere and probably caused by the large proportion of desert sites. Contribution to gamma shows a similar decline with latitude, although the decrease is less steep in the southern hemisphere (Fig. 1a,b). Contribution is positively correlated with alpha diversity^7,33^ (Fig. S1a). In contrast, complementarity shows opposing patterns in the two hemispheres. In the northern hemisphere, complementarity decreases from the equator towards higher latitudes, whereas in the southern hemisphere, the pattern is reversed, resulting in a global exponential increase in endemism from north to south (taxonomic complementarity, R² = 0.91; functional complementarity, R² = 0.91; phylogenetic complementarity, R² = 0.93; Fig. 1a,b, Extended Data Table 1). Complementarity is not correlated with alpha diversity^32^ (Supplementary Information).

**Fig. 1.**
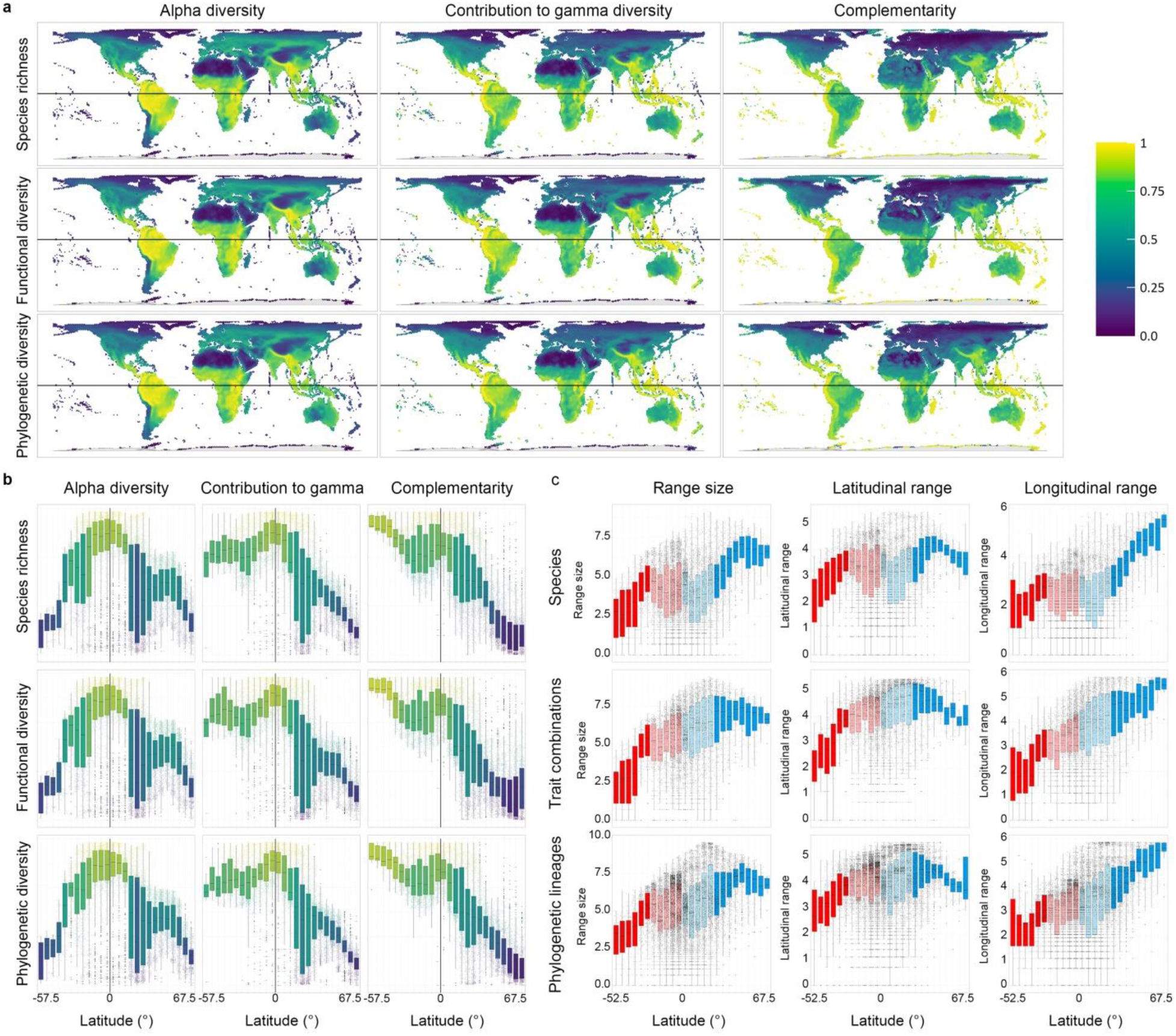
Global patterns of avian diversity, endemism, and range size. **a**, Alpha diversity, contribution to gamma diversity (“weighted endemism”), and complementarity for three facets of diversity (species richness, functional diversity, phylogenetic diversity) mapped across the world (n = 14,640 grid cells). Cells are coloured according to relative rank (1 highest, 1/14,640 lowest values). **b**, Rank alpha diversity, contribution to gamma diversity, and complementarity against latitude shown for three facets of diversity (species richness, functional diversity, phylogenetic diversity). Values (n = 14,640 grid cells) are summarized in latitudinal bands of 5° (n = 26 latitudinal bands). Points are coloured according to relative rank, boxplots according to median values. **c**, Absolute range size, latitudinal range, and longitudinal range (log-transformed) for units representing three facets of diversity (species, functional trait combinations, phylogenetic lineages). Each unit is assigned to a latitudinal band of 5° (n = 25 latitudinal bands) according to the latitudinal midpoint its range. Colours represent southern (red) and northern (blue) hemispheres; latitudinal bands inside the tropics are lighter coloured. In the southern hemisphere, ranges decrease towards higher latitudes; in the northern hemisphere, absolute range size and latitudinal range show a peak at mid-latitudes, and longitudinal range increases towards higher latitudes.

In accordance with the south-north decrease in complementarity, median range size for birds also shows a south-north increase (species, R² = 0.74; functional traits, R² = 0.79; phylogenetic lineages, R² = 0.83), as does latitudinal range size (species, R² = 0.54; functional traits, R² = 0.39; phylogenetic lineages, R² = 0.35), and longitudinal range size (species, R² = 0.72; functional traits, R² = 0.94; phylogenetic lineages, R² = 0.89; Fig. 1c, Extended Data Table 2). If analysed separately by hemisphere, however, latitudinal range size strongly decreases from the equator towards higher latitudes in the southern hemisphere for functional trait combinations (R² = 0.90, p < 0.001) and phylogenetic lineages (R² = 0.88, p < 0.001), but shows a hump-shaped relationship with latitude in the northern hemisphere (functional trait combinations, R² = 0.81, p < 0.001, peak at 27.5 ° N; phylogenetic lineages, R² = 0.79, p < 0.001, peak at 42.5 °N).

By using an endemism measure that is only influenced by the overlap in assemblage composition, but not assemblage richness (alpha diversity), and by including all global areas, we reveal a global north-south gradient in avian endemism with a peak in the southern hemisphere. On average, southern hemisphere bird assemblages show the highest proportion of irreplaceable, range-restricted species, functional trait combinations and phylogenetic lineages. This pattern for endemism measured as complementarity is strikingly different from that for contribution to gamma diversity (i.e. *weighted endemism*), the most-commonly used measure for endemism^13–15^ (Figs. 1a,b, 2). In addition to known diversity hotspots such as the tropical Andes, Madagascar, and New Guinea, we reveal endemism peaks at sites that are not normally considered diversity hotspots, such as oceanic islands, and most notably, the sub-Antarctic islands and the Antarctic continent, i.e. sites that are commonly omitted from large-scale analysis of diversity^8–11^ because of their low alpha diversity and small landmass (Figs. 1a, 2). Their conservation significance therefore owes not only to the shear abundance of birds found on them^43–44^, but also to their global contribution to the retention of the extant avifauna.

**Fig. 2.**
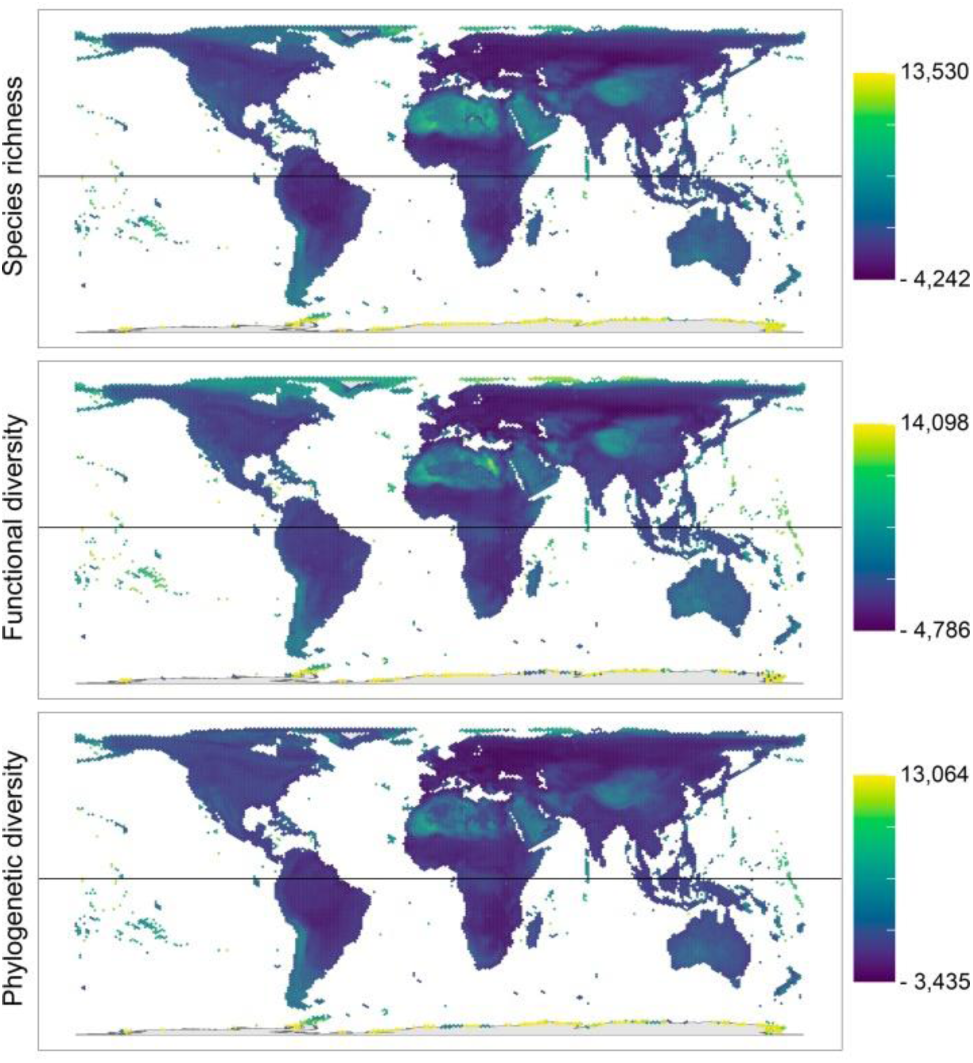
Changes in the relative importance (rank) of sites with regard to their complementarity (“corrected weighted endemism”) vs their contribution to gamma diversity (“weighted endemism”). Grid cells (n =14,640 grid cells) are coloured ranging from maximum increase to maximum decrease in rank differences (rank complementarity – rank contribution to gamma diversity). Antarctica, the sub-Antarctic islands, the Arctic, oceanic islands in the tropics, desert regions, and the southern Andes show the largest increase in rank; temperate regions in the Palaearctic show the largest decrease in rank.

Our results reveal a previously overlooked difference between hemispheres in endemism and range size. To assess the potential significance of these differences for the conservation of northern-hemisphere and southern-hemisphere species, we explored these differences further. First, we determined if the differences in endemism and range size influence patterns of gamma diversity by calculating taxonomic, functional, and phylogenetic gamma diversity for each latitudinal band of grid cells. Since endemism and range size are influenced by the availability of landmass, we also quantified available landmass, approximated by the number of grid cells per latitudinal band, and then calculated density (i.e. gamma diversity/available landmass) for each latitudinal band of grid cells.

Gamma diversity shows similar patterns in both hemispheres, with a peak in the tropics and a decline towards higher latitudes, albeit with a slightly faster decrease outside the tropics in the southern hemisphere (extratropical, species richness, north, R² = 0.98, south, R² = 0.98; functional diversity, north, R² = 0.97, south, R² = 0.99; phylogenetic diversity, north, R² = 0.97, south, R² = 0.98, Fig. 3d). In contrast, available landmass shows strikingly opposing relationships in the two hemispheres, especially outside the tropics (Fig. 3a,b): it continuously increases in the northern hemisphere up until 65 ° N, but it decreases up until 40 ° S in the southern hemisphere (Fig. 3b,c), resulting in an overall continuous increase of available landmass from south to north. This opposing pattern across hemispheres necessarily contradicts the idea^45^ that the increase in gamma diversity from the poles towards the tropics can be explained by larger available landmass and larger ranges in the tropics; absolute range size, latitudinal range size, and longitudinal range size all generally follow the increase in available landmass from south to north (Fig. 1c, Fig. 3b).

**Fig. 3.**
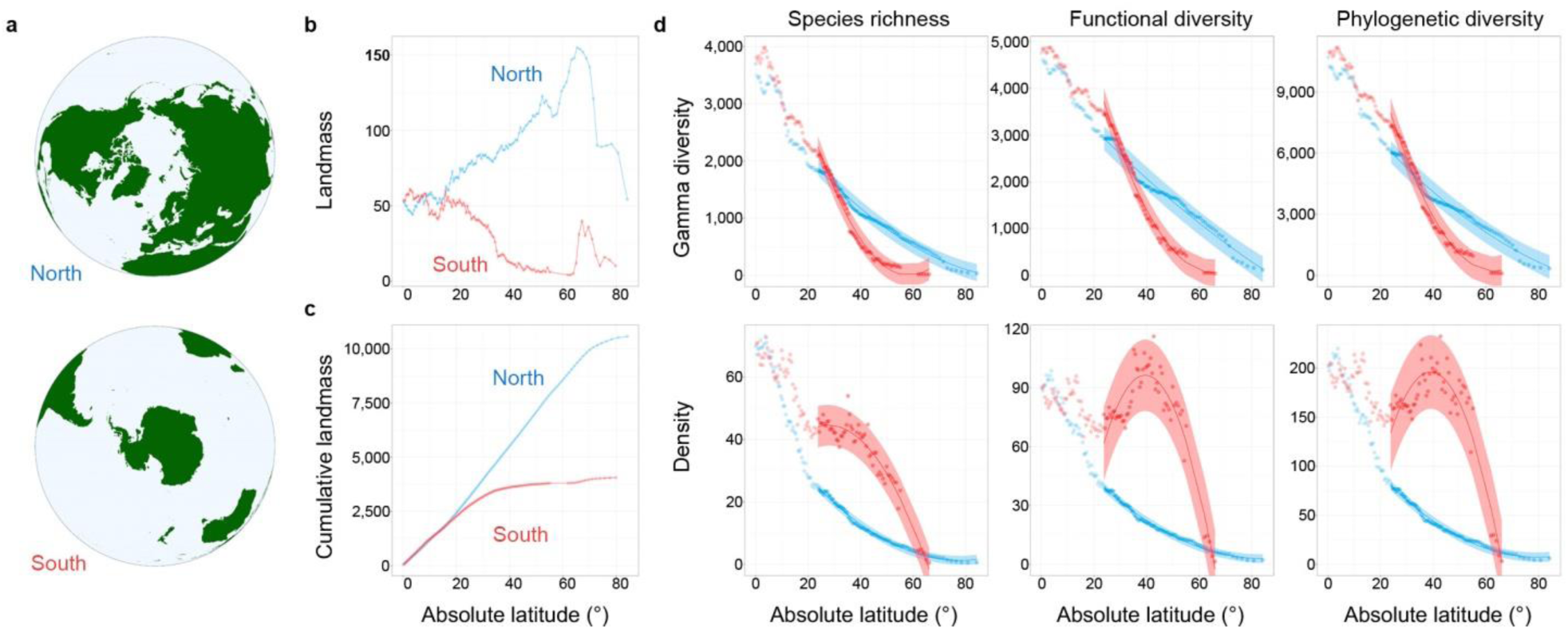
Landmass and gamma diversity in the southern and northern hemisphere. **a**, Extratropical landmasses (green) in the northern hemisphere (above) and southern hemisphere. **b**, Landmass (approximated by number of grid cells) per latitudinal band (n = 252 bands) in the northern (blue) and southern (red) hemisphere against absolute latitude. **c**, Cumulative landmass (number of grid cells) from the equator to the poles in the northern (blue) and southern (red) against absolute latitude. **d**, Gamma diversity (above) and density (gamma diversity/landmass) against absolute latitude in the northern (blue) and southern (red) hemisphere. Gridcells (n = 14,640) are divided into latitudinal bands (n = 245 bands) according to their latitudinal midpoint; gamma diversity is calculated for each latitudinal band. Gamma diversity decreases towards higher latitudes in both hemispheres. Density decreases towards higher latitudes in the northern hemisphere (n = 76 extratropical latitudinal bands); in the southern hemisphere outside the tropics (n = 59 latitudinal bands), density peaks at mid-latitudes. The shading around the regression lines represent 95% confidence intervals.

The uneven distribution of landmass affects latitudinal patterns of density: in the northern hemisphere, density declines continuously from the tropics to the highest latitudes (species richness, R² = 0.99; functional diversity, R² = 0.99; phylogenetic diversity, R² = 0.99; Fig. 3d), whereas in the southern hemisphere, the relationship is hump-shaped outside the tropics, with a peak at mid-latitudes (species richness, R² = 0.94; functional diversity, R² = 0.87; phylogenetic diversity, R² = 0.86; Fig. 3d); for functional and phylogenetic diversity these peaks represent the highest values observed along the entire latitudinal gradient (Fig. 3d). The increase of available landmass from south to north corresponds with the observed increase in range sizes, especially the increase in longitudinal ranges. At the same absolute latitude, species in the northern hemisphere can expand their ranges over a large area of similar climatic conditions^45–47^, whereas in the southern hemisphere, longitudinal ranges are restricted because landmasses are separated by vast expanses of open ocean (Fig. 3a). The smaller range sizes and the higher isolation of landmasses in the southern hemisphere result in reduced overlap in assemblage composition and, consequently, the higher values of contribution to gamma and complementarity observed in the southern hemisphere.

The global north-south increase in endemism markedly departs from current understanding of global diversity patterns. In line with recent studies that showed that southern hemisphere ecosystems, especially the sub-Antarctic and Antarctic, are more diverse than previously thought^48^, we demonstrate that—on average—southern hemisphere assemblages harbour among the highest percentages of irreplaceable, range-restricted species, functional trait combinations and phylogenetic lineages, which makes these assemblages potentially most vulnerable to changes^21,39,49^. Similarly, while latitudinal gradients of diversity are similar in both hemispheres ^40^, the diversity is distributed over a much smaller and more disconnected area in the southern hemisphere, leading to a global peak of functional and phylogenetic diversity per area in the mid-latitudes of the southern hemisphere. At the same time, several areas in the southern hemisphere—including Antarctica, the sub-Antarctic islands, oceanic islands in the tropics, and the southern Andes—were among the sites with the highest discrepancy between complementarity (endemism measured as irreplaceability) and contribution to gamma diversity (i.e. *weighted endemism,* the most commonly used measure for endemism). The classic focus on alpha diversity and weighted endemism to identify biodiversity hotspots hence draws conservation actions away from these highly endemic, but species-poor, assemblages.

The observed hemispheric differences in endemism and available landmass suggest potential differences in the way in which northern and southern species might respond to environmental change. For instance, the smaller area and higher discontinuity of available landmass in the southern hemisphere might affect the ability of southern species to shift their ranges^50^. The majority of landmass in the southern hemisphere consists of the southern tips of continents, isolated by vast expanses of ocean, whereas in the northern hemisphere, vast areas of landmass are connected across latitudes and longitudes^45–47^, and the two largest separated blocks of land (Palearctic and Nearctic) are only separated by the comparatively narrow Bering Strait^51–53^. Moreover, for birds with range limits at the southern tip of continents, the nearest poleward landmasses are the sub-Antarctic islands or the Antarctic continent, which have climatic conditions that make them unsuitable as breeding grounds for most birds. If environmental conditions in the current ranges become unsuitable^25,26^, many southern-hemisphere species might therefore not be able to reach suitable new breeding areas. In addition, due to the isolation of landmasses, it is not uncommon for southern-hemisphere species to have their most-closely related species on a different continent^2^, and local extinctions are therefore less likely to be replaced by the functionally or phylogenetically most-closely related species.

To date, most studies on the potential effect of environmental changes on species assemblages are based on data from the northern hemisphere^54^, and it is not clear in which way species assemblages in the two hemispheres might differ in their responses to climatic changes. Our results demonstrate that, in addition to protecting high-richness areas, conservation actions should be aimed towards protecting the most-highly unique ecosystems that are vulnerable due to their high dependence on endemics, that is, their high dependence on irreplaceable species, functional trait combinations, and evolutionary lineages, and which are most prevalent in the southern hemisphere.

## Methods

### Species distributions

We used the most recent data on avian breeding ranges from the IUCN^37^. We re-projected all ranges into an equal-area projection (EPSG 6933), and then mapped the distribution in a hexagonal grid of 14,640 cells with a size of 11,610 km^2^, roughly equal to 1 ° x 1 ° at the equator, the smallest suitable size for global analyses^55^. For each bird species, we determined range size and the latitudinal midpoint of its distribution. We used latitudes 23.5° N and 23.5° S as cut-offs to divide all grid cells into three groups according to their latitudinal midpoint (north, 7,631 grid cells : tropics, 5,548 grid cells; south, 1,461 grid cells).

### Alpha and gamma diversity

#### Taxonomic diversity

We measured taxonomic alpha diversity as species richness, i.e. the number of bird species found in each site; we measured taxonomic gamma diversity as the total number of bird species.

#### Functional diversity

We measured functional diversity as the diversity of functional-trait combinations of birds. We selected eight morphological traits^56,57^: culmen length, beak length from tip to nares, beak depth, beak width, tarsus length, wing length (carpal joint to wingtip), Kipp’s distance (carpal joint to tip of outermost secondary), and tail length. We obtained trait measurements for all species from AVONET^38^. We used Principal Coordinates Analysis (PCoA) to project all bird species into one common four-dimensional trait space where they were arranged according to the differences in their trait combinations. Since the position in trait space represents a species’ niche position but not its niche size^58,59^, we assigned a volume around each bird species with a radius equivalent to 1/15 of the length of the shortest PCoA axis, which is a conservative approach (using 1/10 lead to very similar results). This approach is similar to calculating functional diversity as cumulative trait probability densities with fixed standard deviation (TPD_c_)^60^, with virtually identical FD values (R^2^ = 0.99–1.0), but it is much faster and facilitates the inclusion of more species and trait dimensions. We measured functional alpha diversity as the volume of trait space occupied by the bird species present in a geographic grid cell, ignoring the overlap between the volumes of individual species as well as any “empty” regions in the trait space^59^. Likewise, we measured functional gamma diversity as the volume of trait space occupied by all bird species from the global dataset.

#### Phylogenetic diversity

We measured phylogenetic alpha and gamma diversity as Faith’s PD^61^, the combined length of all branches that connect the bird species present in a grid cell (alpha), or the all bird species in the dataset (gamma). We obtained 1000 dated phylogenetic trees from birdtree.org^39^ and calculated a consensus tree based on the phylogenetic backbone by ref. 62.

### Endemism

We quantified local endemism with two measures: contribution to gamma diversity (“contribution”) and complementarity. Both are derived from measures for comparing the diversity of species’ functional roles in ecological networks^33,59^ and species’ contribution to functional and phylogenetic diversity^35^, which can also be applied to large-scale comparisons of diversity across communities^36^.

#### Contribution to gamma diversity (weighted endemism)

Contribution to gamma diversity represents the contribution of each site to global gamma diversity^33,35,59^. It is calculated by weighting local alpha diversity according to how often each element (species, functional trait combination, branch in the phylogeny) is found in other sites (i.e., a species found in n sites contributes 1/n to gamma in each of the communities in which it occurs). These local *weighted alpha* diversities add up to gamma; the fraction between *weighted alpha* / gamma represents the contribution to gamma. Contribution to gamma diversity is strongly correlated with local alpha diversity (Supplementary Information, Fig. S1). Complementarity measures the degree to which the elements of the local alpha diversity (species, functional trait combinations, branches in the phylogeny) are shared with other sites^33,36^. It is calculated as *weighted alpha diversity* / alpha diversity^33^. Although derived differently, contribution to gamma and complementarity are conceptually similar to the measures of weighted endemism and corrected weighted endemism^4,31,63,64^, provided that they are measured across sites of equal size (e.g. a grid). Contribution is also equivalent to *range(-size) rarity*^32,65^; complementarity is equivalent to *proportional range rarity*^32^. Since complementarity is independent of local alpha diversity (Supplementary Information, Fig. S1), it is a more suitable measure for the irreplaceability of a site or region than contribution to gamma.

### Relationship with latitude, available landmass

#### Alpha diversity against latitude

We divided all grid cells into latitudinal bands of 5° from 60°S to 70° N (n = 26 bands) and summarized median taxonomic, functional and phylogenetic contribution and complementarity and median range size for each band. We used linear regression models to relate median values to latitude.

#### Gamma diversity against latitude

We divided all grid cells into latitudinal bands according to their latitudinal midpoint (n = 252 bands). We calculated taxonomic, functional, and phylogenetic gamma diversity per band, but we removed six latitudinal bands with fewer than three grid cells. For each band, we also quantified available landmass, approximated by the number of grid cells. We excluded grid cells south of 67°S because only a tiny fraction of their landmass is ice-free, and an inclusion would therefore misrepresent the relationship between gamma diversity and available landmass (an analysis including the cells south of 67° S, however, led to almost identical results). We used linear models to test the relationship between gamma diversity, density (gamma diversity / available landmass) and latitude.

## Data availability

Bird range maps are available from BirdLife International (version 2020.1, http://datazone.birdlife.org)^37^, but restrictions apply to the availability of these data, which were used under license for the current study. Data are, however, available from the authors upon request and with permission of Birdlife International. Trait data for all birds are available from AVONET^38^, bird phylogenies are available from phylotree.org^39^. Data to repeat the analyses and figures will be provided as Supplementary Information.

## Code availability

R code for statistical analyses will be made available as Supplementary Information.

## Acknowledgements

This work was supported by ARC SRIEAS Grant SR200100005 Securing Antarctica’s Environmental Future.

## Author contributions

D.M.D. conceived the study, D.M.D and S.LC. developed the study design; D.M.D. handled all data processing, developed the methods and carried out analyses; D.M.D. and S.L.C. interpreted the results; D.M.D. generated the figures and wrote the first draft; D.M.D. and S.L.C. revised the manuscript.

## Additional information

Supplementary Information is available for this paper.

Correspondence and requests for materials should be addressed to D. M. Dehling.

