## Supplementary Information for "Global increase in the endemism of birds from north to south"

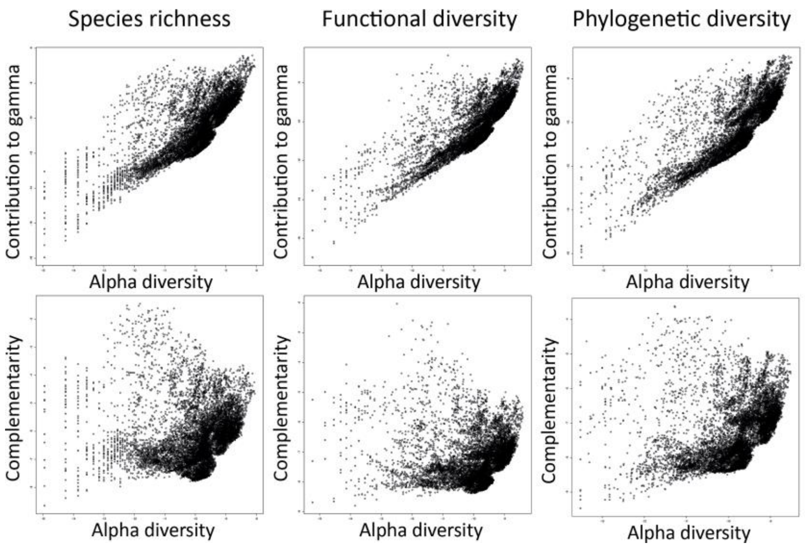


Fig. S1. Relationship (log-log) between local alpha diversity and local contribution to gamma diversity (“weighted endemism”) and complementarity for three facets of diversity (taxonomic, functional, and phylogenetic diversity). Contribution to gamma diversity is strongly correlated with local alpha diversity; complementarity is not correlated with local alpha diversity.

Extended Data Table 1 | Median log complementarity against latitude for three facets of diversity (taxonomic, functional, phylogenetic diversity). n = 26 latitudinal bands

| | Taxonomic complementary | | | |
|---|---|---|---|---|
| | Estimate | t | $R^2$ | p-value |
| Latitude | -0.09 ± 0.006 | -15.8 | 0.91 | < 0.001 |
| | Functional complementarity | | | |
| | Estimate | t | $R^2$ | p-value |
| Latitude | -0.08 ± 0.005 | -15.6 | 0.91 | < 0.001 |
| | Phylogenetic complementarity | | | |
| | Estimate | T | $R^2$ | p-value |
| Latitude | -0.08 ± 0.005 | -18.3 | 0.93 | < 0.001 |

These models estimate the relationship between median complementarity (log-transformed) and latitude. The analysis was conducted across 14,640 grid cells, summarized in latitudinal bands of 5° (n = 26 latitudinal bands). The p-values are computed using two-sided tests.

Extended Data Table 2 | Median range size (absolute range size, latitudinal range size, longitudinal range size) against latitude for units that represent avian taxonomic, functional, and phylogenetic diversity (species, functional-trait combinations, phylogenetic lineages, respectively). n = 26 latitudinal bands

*Species*

| | Latitude | | | |
|---|---|---|---|---|
| | Estimate | t | $R^2$ | p-value |
| Absolute range size | 0.16 ± 0.02 | 8.16 | 0.74 | < 0.001 |
| Latitudinal range size | 0.05 ± 0.01 | 5.18 | 0.54 | < 0.001 |
| Longitudinal range size | 0.12 ± 0.02 | 7.82 | 0.72 | < 0.001 |

*Functional-trait combinations*

| | Latitude | | | |
|---|---|---|---|---|
| | Estimate | t | $R^2$ | p-value |
| Absolute range size | 0.19 ± 0.02 | 9.17 | 0.79 | < 0.001 |
| Latitudinal range size | 0.06 ± 0.02 | 3.83 | 0.39 | < 0.001 |
| Longitudinal range size | 0.15 ± 0.01 | 18.7 | 0.94 | < 0.001 |

*Phylogenetic lineages*

| | Latitude | | | |
|---|---|---|---|---|
| | Estimate | t | $R^2$ | p-value |
| Absolute range size | 0.15 ± 0.01 | 10.6 | 0.83 | < 0.001 |
| Latitudinal range size | 0.04 ± 0.01 | 3.49 | 0.35 | 0.002 |
| Longitudinal range size | 0.12 ± 0.01 | 13.7 | 0.89 | < 0.001 |

These models estimate the relationship between different measures of median range size and latitude. All p-values are computed using two-sided tests.
